# Migration of Cytotoxic T Lymphocytes in 3D Collagen Matrices

**DOI:** 10.1101/2020.01.14.906016

**Authors:** Z. Sadjadi, R. Zhao, M. Hoth, B. Qu, H. Rieger

## Abstract

CD8^+^ cytotoxic T lymphocytes (CTL) and natural killer (NK) cells are the main cytotoxic killer cells of the human body to eliminate pathogen-infected or tumorigenic cells (= target cells). To find their targets they have to navigate and migrate through a complex biological microenvironments, a key component of which is the extracellular matrix (ECM). The mechanisms underlying killer cell’s navigation are not well understood. To mimic an ECM we use a matrix formed by different collagen concentrations, and analyze migration trajectories of primary human CTLs. Different migration patterns are observed and can be grouped into three motility types: slow, fast and mixed. The dynamics are well described by a two-state persistent random walk model which allows cells to switch between slow motion with low persistence, and fast motion with high persistence. We hypothesize that the slow motility mode describes CTLs creating channels through the collagen matrix by deforming and tearing apart collagen fibers, and that the fast motility mode describes CTLs moving within these channels. Experimental evidence supporting this scenario is presented by visualizing migrating T cells following each other on exactly the same track and showing cells moving quickly in channel-like cavities within the surrounding collagen matrix. Consequently, the efficiency of the stochastic search process of CTLs in the ECM should strongly be influenced by a dynamically changing channel network produced by the killer cells themselves.

## I. INTRODUCTION

Cytotoxic T lymphocytes (CTLs) are fully activated CD8^+^ T cells, which are key players of the adaptive immune system to eliminate tumorigenic or pathogen-infected cells [1]. CTLs need to find the cognate antigens presented by target cells, for example pathogen-infected or tumorigenic cells, in order to eliminate those aberrant cells in the immunosurveillance process. These target cells are often low in number in the early stages of disease development [2–4]. Thus, the ability of CTLs to efficiently navigate and search is crucial for an efficient immune response. Migration behavior of immune cells in the body and the search strategies they might follow is currently of great interest in physics and biology [5–7]. Migration of naive T cells in lymph nodes reportedly follows a Brownian or even subdiffusive dynamics [8–10], but switchings between fast and slow motility modes have also been observed [11]. Outside the lymph node, activated T cells are destined to find their targets in peripheral tissues, most of which are characterized by dense extracellular matrix (ECM) [12]. Here a faster migration, e.g. via longer phases of superdiffusive dynamics or less switchings to the slow diffusive mode, is advantageous to scan a larger tissue efficiently. For instance, it was reported that the dynamics of CD8^+^ T cells in infected brain tissue resembles a Levy walk [13].

The extracellular matrix (ECM)– the major component of peripheral tissues– mainly consists of collagens and has essential regulatory roles in nearly all cellular functions. Some collagens have inhibitory effects on the function of different immune cells [14, 15]. In various types of cancer, the collagen network becomes dense, stiff and linearized in the vicinity of tumors, facilitating the transport of cancerous cells and making the ECM an important player in cancer metastasis, intravasion and prognosis [16–21]. Additionally, the proliferation of CTLs is impaired in a high-density collagen matrix [22]. Recently, different immune cells have been investigated in immunotherapy studies as potential drug delivery vehicles into tumors [23, 24]. Understanding the migration and interactions of immune cells in collagen networks is crucial to unravel the underlying details of the immune response and design effective treatment strategies.

Collagen-based assays have been used to investigate the migration of lymphocytes in ECM and study the possible underlying mechanisms of immune interactions with ECM [14, 25–29]. In a recent study, collagen hydrogels were employed to compare migration patterns of human CD8^+^ T cells in aligned and nonaligned collagen fibers microenvironments, resembling tumor cells and normal tissues, respectively [30].

In this study, we use bovine collagen to construct a 3D environment in vitro as a model for the ECM. The trajectories of primary human CTLs in collagen matrices with different concentrations are analyzed for two blood donors. We find three different types of motion in both donors: The migration of CTLs can be categorized into slow, fast and mixed sub-groups and show that a persistent random walk model with two different motility states, a slow one and a fast one, and transitions between describes the experimental data accurately. Finally we provide a biophysical interpretation of the two motility modes of T cells in collagen matrix related to channel formation and movement within channels.

## II. MATERIALS AND METHODS

Ethical considerations: Research carried out for this study with human material (leukocyte reduction system chambers from humanblood donors) is authorized by the local ethic committee (declaration from 16.4.2015 (84/15; Prof. Dr. Rettig-Stürmer)).

Human primary cytotoxic T lymphocytes (CTL) isolation, stimulation and nucleofection of CTLs: Peripheral blood mononuclear cells (PBMCs) were obtained from healthy donors as previously described [33]. Human primary CTLs were negatively isolated from PBMCs using Dynabeads^*TM*^ Untouched^*TM*^ Human CD8 T Cells Kit (ThermoFisher Scientific) or CD8^+^ T Cell Isolation Kit, human (Miltenyi Biotec), stimulated with Dynabeads^*TM*^ Human T-Activator CD3/CD28 (ThermoFisher Scientific) in AIMV medium (ThermoFisher Scientific) with 10% FCS and 33 U/mL of recombinant human IL-2 (ThermoFisher Scientific). 48 hours after stimulation beads were removed and 5 × 10^6^ CTLs were electroporated with 2 *μg* plasmid (H2B-GFP to label nucleus [34] or pMax-mCherry to label cell bodies) using the Human T cell nucleofector kit (Lonza). Medium was changed 6h after nucleofection and transfected CTLs were maintained in AIMV medium (ThermoFisher Scientific) with 10% FCS and 33 U/mL of recombinant human IL-2 (ThermoFisher Scientific). Cells were used 24-36 hours after nucleofection [35].

### A. 3D live cell imaging with lightsheet microscopy

3D live cell imaging using lightsheet microscopy was conducted mainly as described previously [36]. Briefly, human primary CTLs were resuspended first in PBS (ThermoFisher Scientific), afterwards neutralized collagen stock solution (bovine collagen type I, 8 mg/mL, Advanced Biomatrix) was added to a final concentration of 2 mg/ml, 4 mg/ml, or 5 mg/ml collagen with a cell density of 10 × 10^6^ cells/ml. 20 *μl* of this cell/collagen mixture was loaded in a capillary. The capillary was closed and incubated for 60 min in an incubator. Afterwards, the polymerized collagen rod was pushed out hanging in the medium at 37°*C* with 5% CO_2_ for equilibration for another 60 min. To visualize collagen structure, analyzed collagen matrix was stained with 50 *μ*g/ml Atto 488 NHS ester (ThermoFisher Scientific) in AIMV medium at room temperature after collagen polymerization. Afterwards, the matrix was washed in AIMV medium. After collagen polymerization, cells in the matrix with or without collagen staining were incubated in AIMV medium with 10% FCS at 37°*C* with 5% CO_2_ for 1 hour. Afterwards, the migration of cells was visualized by lightsheet micoscopy (20×objective) at 37°*C* with a z-step size of 1 *μm* and a time interval of 30 seconds. The migration trajectories were tracked and analyzed using Imaris 8.1.2 (containing Imaris, ImarisTrack, Imaris-MeasurementPro, ImarisVantage; Bitplane AG, software available at http://bitplane.com) [36].

### B. Data Analysis

The experimental trajectories consist of a set of T cell positions recorded after equal time intervals. Every two successive recorded positions are used to calculate the instantaneous velocity, and every three of them to extract the corresponding turning angle *ϕ*. The values of *ϕ* around zero represent a tendency to continue along the previous direction of motion, i.e. a persistent motion. In contrast, *ϕ* values close to *π* indicate reversing the direction of motion. Accordingly we define the instantaneous persistence as 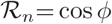 and the average persistence as 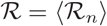.

We apply a minimum duration threshold of 6 frames (i.e. 3 mins) to filter out short trajectories, which leads to smooth velocity and turning-angle distributions. We checked that moderate changes of the minimum duration of trajectories has a negligible influence on the relative population of different categories of T cell migration pattern. Moreover, a second threshold is applied on the duration of trajectories when analyzing the switching statistics between the sub-states of the mixed migration type. In order to minimize the effects of the tracking time window, we only analyze trajectories with the duration of 50 min (i.e. 100 frames). The longest possible tracking time is 60 min since the duration of our live cell imaging is 1 hour. We checked that the results are not significantly affected by a slight reduction of the minimum trajectory duration.

T cells enter the camera field at different times. We shift the starting time of all trajectories, so that all tracks start at the same time (*t*=0). Throughout the manuscript, we use the notation *t* for the time from the beginning of each track and Δ*t* for the time interval between two successive recorded positions.

## III. RESULTS

To investigate migration patterns of CTLs in a 3D environment, we embedded primary human CTLs into collagen matrices and visualized their movements using lightsheet microscopy (Fig. 1). Different concentrations of collagen mimic the ECM of normal tissue (2 mg/ml), soft solid tumor (4 mg/ml) and hard solid tumor (5 mg/ml), respectively [37–39]. The resulting parameters are summarized in Table I. The cross correlation between velocity and persistence, 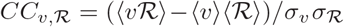, which shows how these two parameters are related, also shows no systematic dependence on the collagen density. We find that 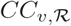 is always positive, which means that faster T cells move more persistently than slower ones. The distributions of velocity, turning-angle and persistence are shown in Fig. 2. The average velocity is higher at lower densities of collagen as expected. The distributions of 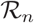 and *ϕ* show a tendency to turn with an angle around 0.4 to 0.5 radian (corresponding to a persistence around 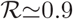).

**FIG. 1.**
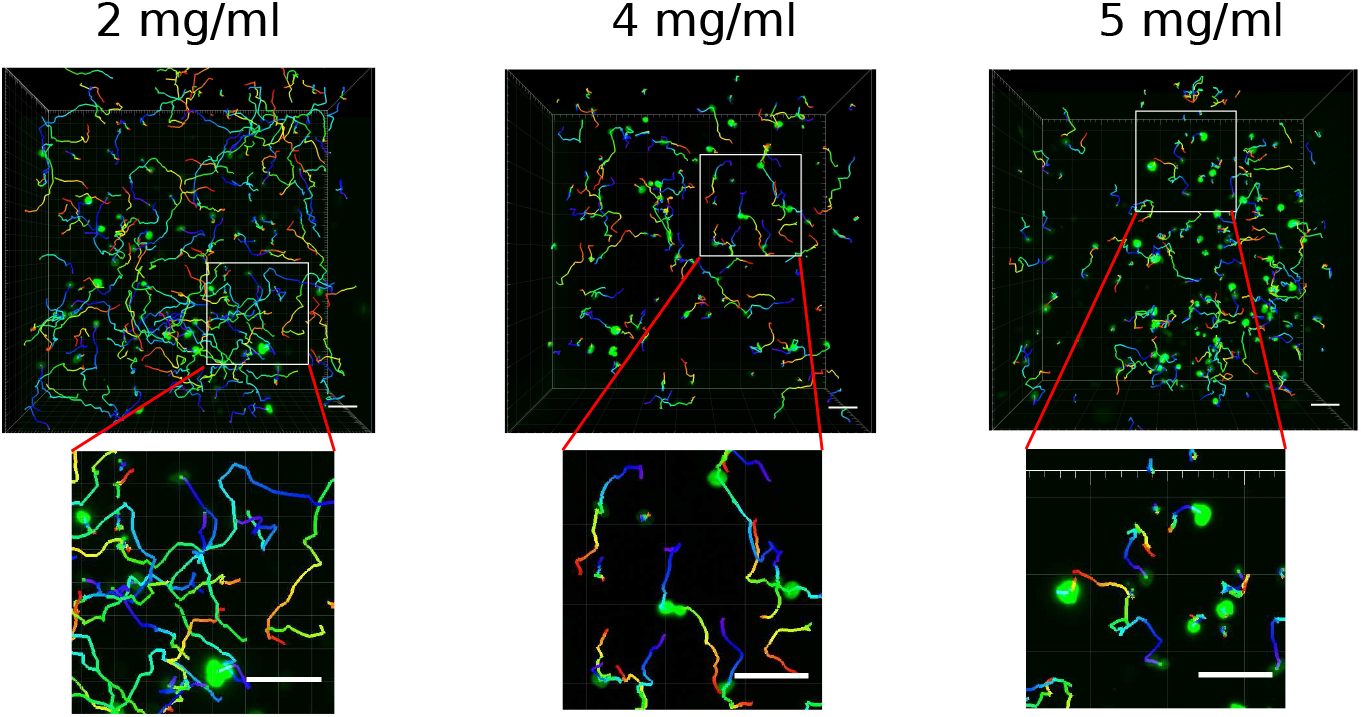
CTL migration trajectories measured by lightsheet microscopy in in 3D collagen matrices with different concentrations. The nuclei of human primary CTLs were labeled with overexpressed Histone 2B-GFP (green). CTL migration trajectories were tracked automatically using Imaris. Scale bars are 40 *μm*. In the lower panels, magnified images of the area marked by the solid squares are shown.

**TABLE I.** Key statistical parameters of T cells in collagen matrices with different densities.

| density ($\frac{\text{mg}}{\text{ml}}$) | Donor 1 | | | Donor 2 | | |
| --- | --- | --- | --- | --- | --- | --- |
|  | 2 | 4 | 5 | 2 | 4 | 5 |
| velocity $v$ ($\frac{\mu\text{m}}{\text{s}}$) | $0.10 \pm 0.002$ | $0.06 \pm 0.002$ | $0.05 \pm 0.003$ | $0.07 \pm 0.004$ | $0.04 \pm 0.002$ | $0.03 \pm 0.001$ |
| persistence $\mathcal{R}$ | $0.30 \pm 0.018$ | $0.36 \pm 0.024$ | $0.35 \pm 0.018$ | $0.30 \pm 0.044$ | $0.36 \pm 0.026$ | $0.44 \pm 0.03$ |
| $CC_{v,\mathcal{R}}$ | 0.64 | 0.35 | 0.35 | 0.36 | 0.29 | 0.31 |

**FIG. 2.**
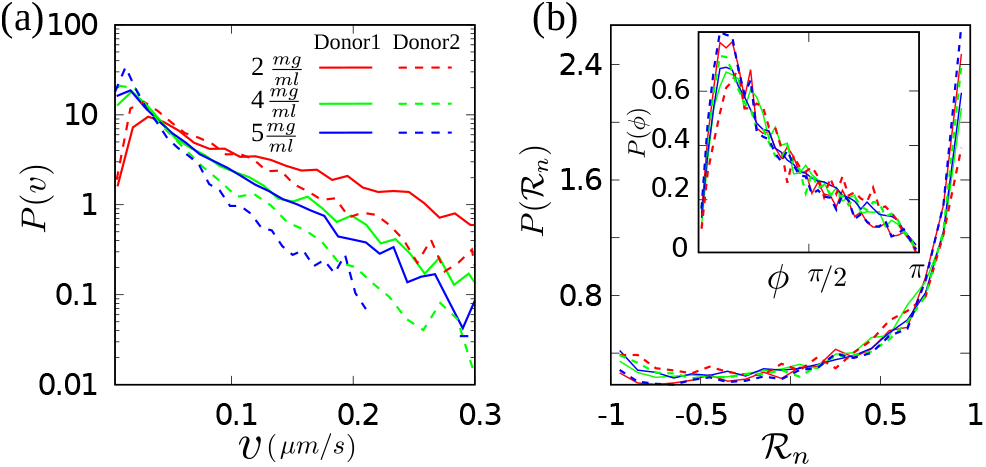
Distributions of (a) velocity, and (b) persistence of T cells in collagen networks with different concentrations. Turning-angle distributions are shown in the inset.

*T cell dynamics.* To better understand the dynamics of T cells in matrices with different collagen concentrations we analyze the velocity autocorrelation *C*_*v*,*v*_ as well as the mean square displacement (MSD) separately for each experimental condition in Fig. 3. The gray dashed line in Fig. 3(b) corresponds to Brownian diffusion. The smaller slope of the MSD curves under all conditions shows that the cell motion is slower than normal diffusion and eventually crosses over to diffusive dynamics at long times. Both the decay of *C*_*v*,*v*_ and the crossover of the MSD to asymptotic diffusion show that the cell orientation is randomized after a while.

**FIG. 3.**
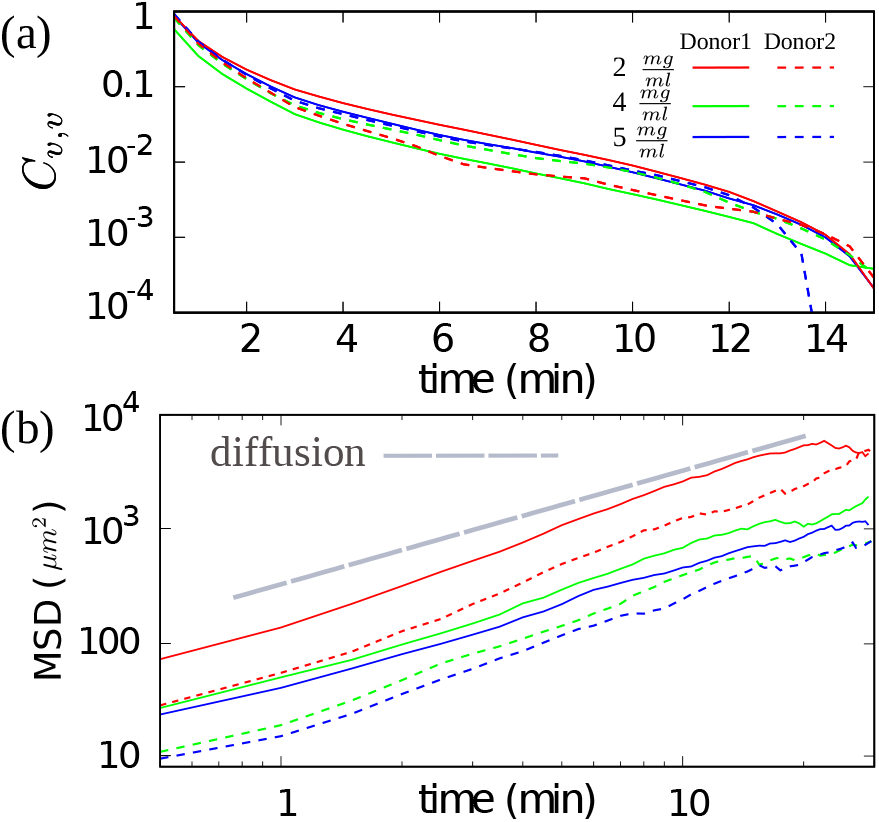
(a) Velocity autocorrelation of CTLs for different donors and collagen concentrations. (b) Time evolution of the mean square displacement of CTLs. The colors and types of the lines are the same as in panel (a).

### A. Three motility groups can be distinguished in CTL dynamics

Single track analysis of CTL trajectories reveals that there are three different types of CTL trajectories: (i) slow T cells which perform a sub-diffusive motion with velocities that always remain below a threshold value, (ii) a faster group with velocities always above a threshold value, and (iii) the third group with velocities which switch between fast and slow modes. Both donors have CTLs of the three types, though with different proportions (shown in Table II). The velocity evolution of typical tracks and a few trajectories for each cell motility type are shown in Fig. 4.

**TABLE II.** Percentage of different motility types of CTLs and the threshold velocities *v*_*c*1_ and *v*_*c*2_ to categorize T cell migration patterns for each experiment with a different donor or collagen density.

| density ($\frac{mg}{ml}$) | Donor 1 | | | Donor 2 | | |
| --- | --- | --- | --- | --- | --- | --- |
|  | 2 | 4 | 5 | 2 | 4 | 5 |
| slow % | 33 | 43 | 48 | 15 | 27 | 26 |
| fast % | 35 | 14 | 19 | 30 | 25 | 24 |
| mixed % | 26 | 37 | 25 | 48 | 40 | 43 |
| $v_{c1}(\mu m/s)$ | 0.075 | 0.055 | 0.04 | 0.05 | 0.02 | 0.02 |
| $v_{c2}(\mu m/s)$ | 0.11 | 0.09 | 0.10 | 0.08 | 0.04 | 0.04 |

**FIG. 4.**
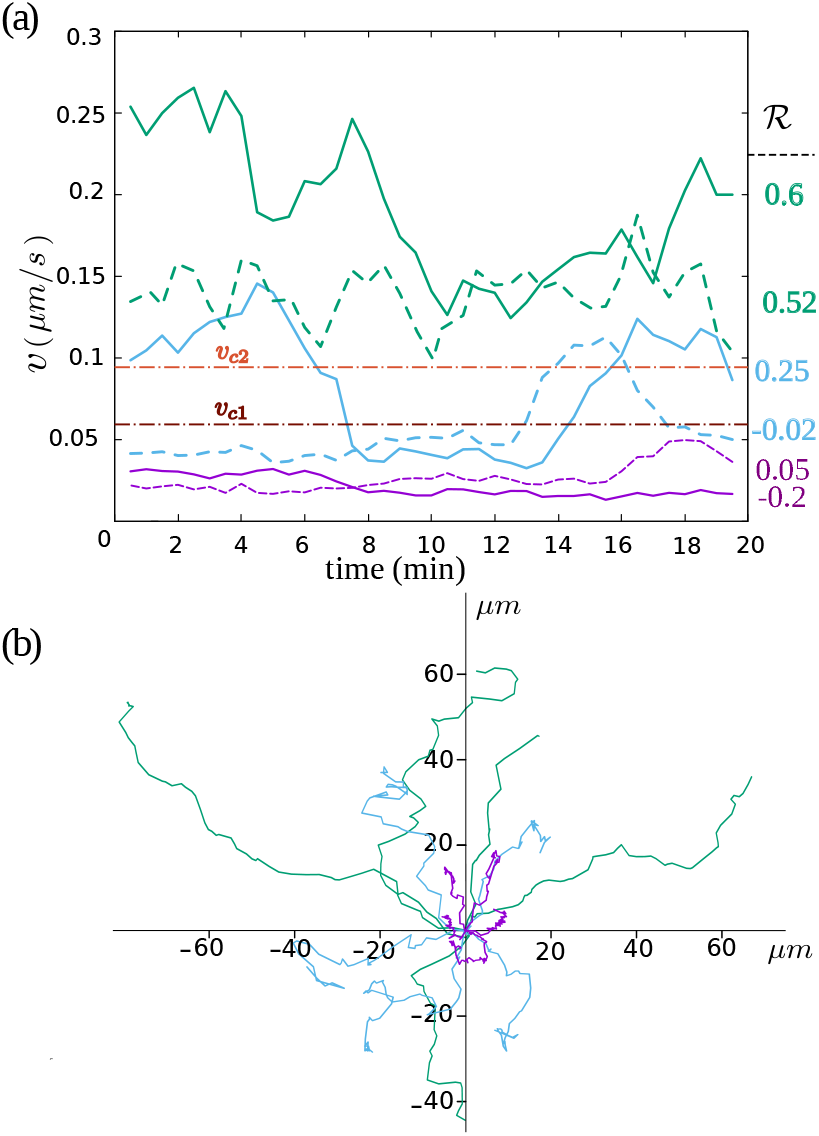
(a) Typical velocities of CTL trajectories. Green, purple and blue colors correspond to a few examplary trajectories of fast, slow and mixed types of motility. The corresponding persistence value 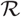 for each trajectory is given on the right *y*-axis. The horizontal lines represent the threshold velocities *v*_*c*1_ and *v*_*c*2_ to categorize T cell migration patterns. (b) Typical trajectories of different motility types. The trajectories belong to donor 1 in the collagen matrix with 2 mg/ml concentration.

We define the three classes of trajectory types - fast, slow, mixed - by introducing two threshold velocities *v*_*c*1_ and *v*_*c*2_ (*v*_*c*2_ > *v*_*c*1_ > 0). A trajectory is classified into the “fast” type if the velocity is at all time *t* larger than *v*_*c*2_, i.e. *v*(*t*) > *v*_*c*2_ for all *t*. It is of the “slow” type if its velocity is always smaller than *v*_*c*1_, i.e. *v*(*t*) < *v*_*c*1_ for all *t* and of the “mixed” type if *v*(*t*) > *v*_*c*2_ for some *t* and *v*(*t*′) < *v*_*c*1_ for some other *t*′. For each experimental data set, distinguished by donor and collagen density, we adapt (*v*_*c*1_*, v*_*c*2_) such that the number of trajectories that does not belong into one of the defined classes (e.g. those with *v*(*t*) > *v*_*c*1_ for all *t* but not *v*(*t*) > *v*_*c*2_ for all *t*) becomes minimal. We checked that in all experiments, the trajectories which do not belong to any of the three identified migration categories remain below 6−8% of all trajectories, when adopting an optimal set of (*v*_*c*1_, *v*_*c*2_). The relative populations of the cells in three migration types moderately change upon varying the threshold velocities *v*_*c*1_ and *v*_*c*2_ around their optimal choices. Throughout the paper, we refer to the fast and slow migration types with *F* and *S* subscripts, and represent the slow and fast substates of the mixed type of migration with subscripts I and II, respectively.

To further confirm that the three populations are not donor-dependent, we pooled data from two donors and analyzed the average velocities and persistence of all populations. Figure 5 (a,b) summarizes the average velocities and persistences of different types in different collagen concentrations for both donors. One observes a moderate increase of the average persistence and a moderate decrease of the average velocity is observed with increasing collagen density. The scatter plots of instantaneous persistence versus velocity, which are shown in Fig. 5(c) in different collagen densities, indicate once again that the faster T cells are more persistent than the slow ones. The MSD of different types of motion are clearly distinguishable (see e.g. Fig. 5(d)). The similarities in the overall time evolution of the MSD curves indicate that the underlying structures guiding the cells are similar. The differences in the level of MSD curves reflect the differences in average velocity of CTLs in various migration types. In the next section, we compare the MSD T cell trajectories with the prediction of a two state random velocity model.

**FIG. 5.**
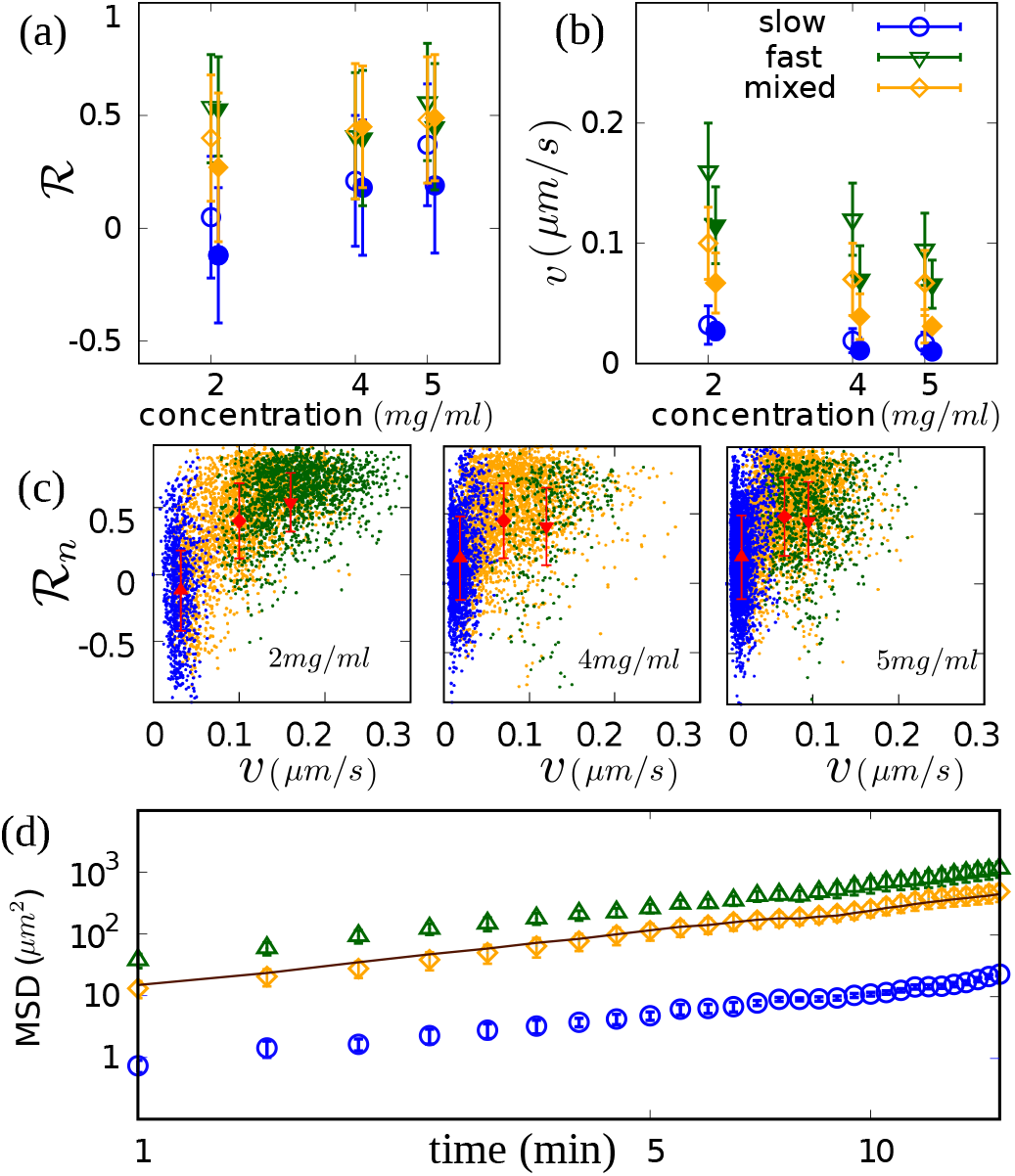
(a) 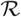 and (b) velocity and their standard deviations for different motility types in two donors represented by open and full symbols. (c) Scatter plots of 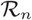 vs *v* in different collagen densities in donor 1, different colors correspond to different migration types as indicated in the legend of part (b). (d) Mean square displacement of different motility types of donor 2 in collagen concentration 5 mg/ml. The solid line represents the MSD of all T cells.

### B. The two-state motility type

In the following we study the mixed-velocity trajectories of T cells in more details.

#### Mean square displacement

The MSD of the mixed type coincides with the total MSD, nearly in all cases (see the solid line in Fig. 5(d)). This shows that the mean velocity of all T cells is nearly the same as the mean velocity of the mixed type when the resident times in the sub-states of the mixed type are taken into account.

#### Exponential distribution of sojourn time in each state

The distribution of the times that the T cells remain in one state before they switch, the so-called sojourn times, follow an exponential decay as shown in Fig. 6. In this example the sojourn time distribution of T cells in the different states of the mixed T cell migration type of two donors is plotted. The exponential decay of these distributions indicates that the transition probabilities are time-independent.

**FIG. 6.**
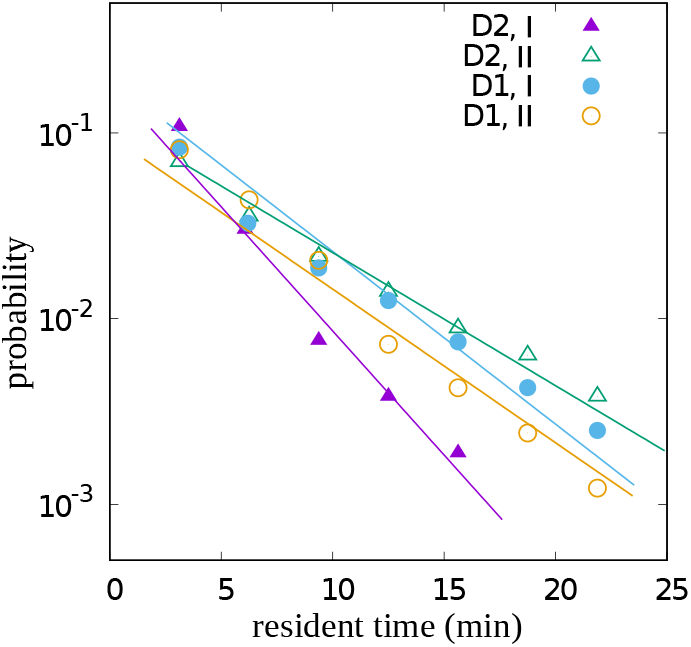
Sojourn time distributions in states I and II of mixed type of T cells in collagen with 4 mg/ml concentration. The lines are the corresponding theoretical estimate for each case (same color). D stands for donor.

#### Probability distribution of persistence in different migration types

Figure 7 shows the probability distribution of the instantaneous persistence of the three different cell types. While the distributions for fast and mixed types show a persistent motion for all collagen concentrations, slow T cells perform anti-persistent motion in 2 mg/ml collagen and become persistent in denser ones. A possible explanation is that the average pore size increases with decreasing collagen density [40] which implies that for low collagen densities, T cells can more easily find some pores around them that are large enough to get into them; this might allow slow T cells to change their direction when they face a constriction while creating a channel. This is, however, less probable in denser collagens, as most pores are smaller than T cells and need equal effort to pass.

**FIG. 7.**
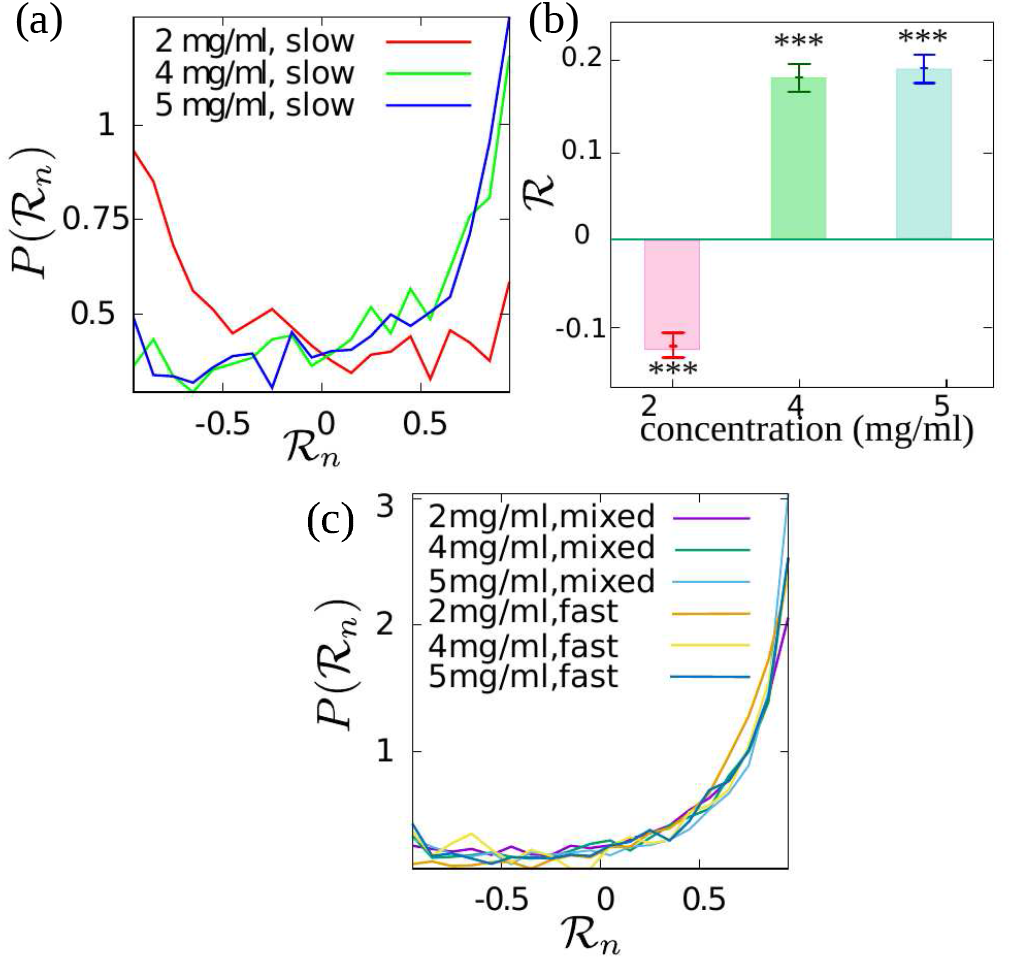
(a) Probability distributions of persistence 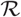 at different collagen concentrations for the slow cell type. (b) Mean persistence 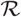 for slow cells (***, *P* < 0.001, t test). (c) Similar to panel (a) for fast and mixed cell types.

### C. CTLs enter channels in the collagen matrix

Next we explored in detail how CTLs migrate in collagen matrices. We fluorescently stained collagen and visualized the movements of CTLs using lightsheet microscopy. We found that during migration, CTLs could enter channels in collagen matrix. Inside the channel they had a high speed which was significantly slowed down when leaving the channel (Fig. 8 (a,c), Suppl. Movie 1). Slow CTLs appeared to be trapped in some channels and moved slowly (Fig. 8(b,c), Suppl. Movie 2). In addition, we observed that in some cases, after one CTL migrated through the matrix, a second CTL followed the same path, indicating that CTLs use channels created by other cells. (see Fig. 8(d), Suppl. Movie 3). When the trajectories of two CTLs overlap the following cell probably moves within the channel created by the leading cell and resulting in a substantial velocity increase of the following cells as shown in Fig. 8(e).

**FIG. 8.**
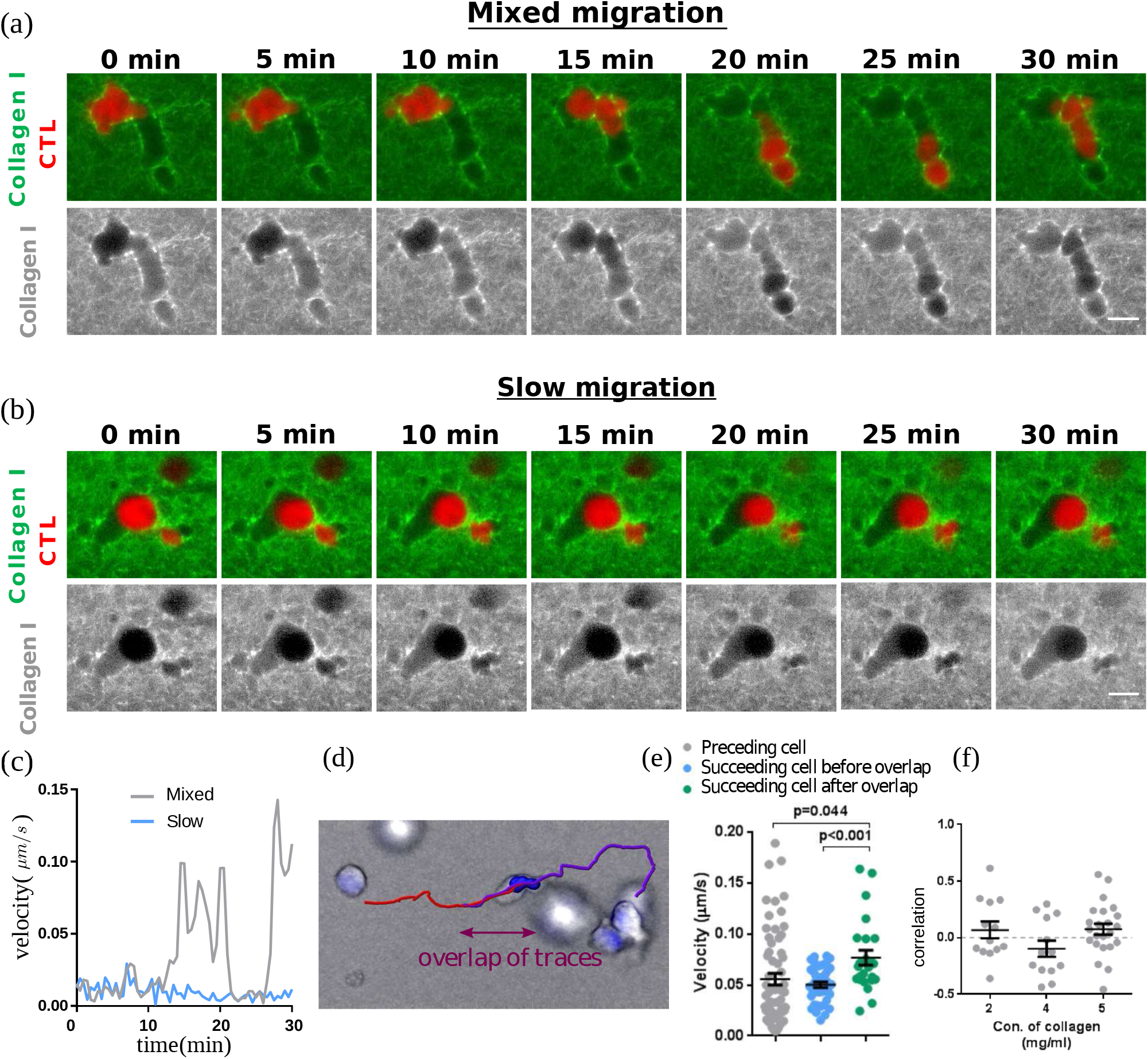
Visualization of CTL migration in a collagen matrix. Primary Human CTLs were transiently transfected with mCherry (red). Coll agen (5 mg/ml) was stained with Atto 488 NHS ester (green or gray as indicated). CTL migration was visualized at 37°C using lightsheet microscopy. Exemplary cells for mixed and slow migration are shown in a and b, respectively (Movies in Suppl. info). One layer of z-stack is presented. c, Quantification of migration velocity at all time points examined. CTL migration trajectories were tracked and analyzed by Imaris. Scale bars are 10 *μm*. (d) An exemplary migrating CTL following another cell in same trace. CTLs were visualized in bright field. Nuclei were labeled with Hoechst 33342 and tacked with imaris. The trajectory of preceding and succeeding cells are red and purple, respectively. Scale bar is 7 *μ*m. (e) The velocity of the preceding cell is similar with the succeeding cell before the overlapping part of traces. When the traces overlap, the velocity of the succeeding cell increases. Each dot presents one time point for the same cell. (f) The correlation of cell body volume and velocity during cell migration. Each dot presents one cell. All results were from at least three independent experiments.

### D. CTLs form channels in the matrix during migration

In order to explore in detail how CTLs migrate in collagen matrices, we investigated whether CTLs can actively create channels while migrating through the matrix. As mentioned earlier, CTLs transiently transfected with red fluorescent protein mCherry were embedded in the fluorescently labeled collagen matrix. As shown in Suppl Movie 4, using light-sheet microscopy, we observed that during migration, CTLs formed protrusions (blob-like structures) at the leading edge, and that these protrusions preferably extended to deformable parts of the matrix and push the matrix aside. After CTLs went through the area, the matrix sprang back to some extent but did not relax to the original form. Therefore, we conclude that through migration, CTLs broaden more easily deformable parts of the matrix to create channels, which plausibly facilitates the migration of the other CTLs entering the same area.

### E. CTL size is not involved in migration speed

To examine whether cell size is involved in migration speed, we analyzed the correlation between cell body volume and velocity for the same cells over time and found no correlation between cell size and migration speed [Fig. 8(f)], indicating that the cell size per se unlikely plays a role in determination of fast, mixed or slow migration types.

## IV. TWO-STATE PERSISTENT RANDOM WALK MODEL

In the following, we show that the experimentally measured T cell trajectories are well described by a stochastic process that involves a persistent random walk with two different motility states [31]. Similar stochastic two-state models have been widely used to describe altering phases of motion in other systems [32].

We adopt a discrete-time approach, since it is best adapted to our experimental data, which consists of the positions of the CTLs at equidistant times. First we focus on the trajectories of the mixed type which involves the slow and the fast motility mode and transitions between them. Later we show that the trajectories of the fast type and those of the slow type are described by the same stochastic process as the mixed type but without transitions from the fast to the slow mode or from low to fast, respectively. This observation complies with the physical interpretation that the slow motility mode is caused by CTLs creating new channels in the collagen matrix and the fast motility mode by CTLs using already existing channels. A persistent random walk in discrete time is a stochastic process for the position of a particle that moves during a time interval Δ*t* with a certain velocity *v* in a certain direction *ϕ*. At the end of the time interval a transition takes place to a new velocity and a new direction. These transitions are characterized by a velocity distribution *F*(*v*) and a turning angle distribution *R*(*ϕ*). A persistent random walk with two motility states involves two velocity distributions, *F*_I_(*v*) and *F*_II_(*v*), and two turning angle distribution *R*_I_(*ϕ*) and *R*_II_(*ϕ*), for the slow (I) and fast (II) motility mode, and transitions between the two motility states characterized by transition probabilities *κ*_I→II_ and *κ*_II→I_ for switching from state I to II and vice versa. These probabilities are estimated by the inverse of the sojourn time in the two states of mixed trajectories, e.g. 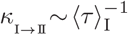. ConIstant probability transitions *κ*_I→II_ (*κ*_II→I_) lead to an exponential distribution of the sojourn time 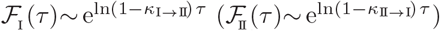. As a first approximation, 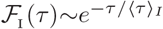 and 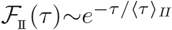 are plotted in Fig. 6, which show a very good agreement with the experimental resident time distribution in each state of the mixed motion.Introducing the probability density functions 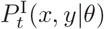 and 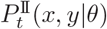 for the probability to find the walker at position (*x, y*) along the direction *θ* at time *t* in each of the motility states, the temporal evolution of the stochastic process can be described by the following set of coupled master equations:

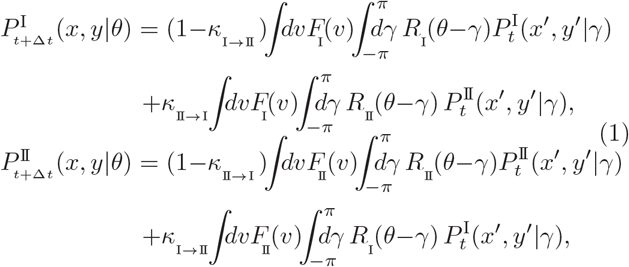

with *x*′ = *x*−*v*Δ*t* cos *θ* and *y*′ = *y*−*v*Δ*t* sin *θ*. By solving these sets of master equations, one can evaluate arbitrary moments of the position of the walker, such as the mean square displacement (MSD). The analytical details to calculate the MSD are presented in the appendix. It should be emphasized that the derived formula for the MSD, i.e. the 2nd moment of the position, depends only on the first two moments of the velocity distributions *F*_I_(*v*) and *F*_II_(*v*), and the first moment of the turning angle distribution.

We extracted the model parameters from the experimental data analysis, thus, there remains no free parameter to be tuned. In case of the mixed type, moments of velocity (〈*v*〉_I_, 〈*v*〉_II_, 〈*v*^2^〉_I_ and 〈*v*^2^〉_II_,〈*v*〉_II_) in each state, are calculated by averaging over local velocities of trajectories. The persistencies 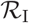 and 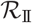 are measured by averaging over *cosine*s of turning angles (see Eq.3). The transition probabilities *κ*_I→II_ and *κ*_II→I_ are the inverse of average sojourn time in states I and II, respectively. In case of the fast and slow types, as explained in the Appendix, the model shrinks to a one state model. The parameters〈*v*〉_j_, 〈*v*^2^〉_j_ and 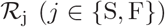. All extracted model parameters are summarized in Tables III and IV for mixed state and fast/slow states, respectively. Note that according to our initial hypothesis the model parameters for the stochastic process describing the trajectories of the slow type (S) should be identical to the model parameters with index I of the mixed type and for the fast type identical to the ones with index II. By comparing the corresponding values in tables III and IV one observes that they do approximately agree, except for the slow motility type where significant differences between the (S) and the (I) parameters occur. We attribute this difference to variations in the local environment in which CTLs move: CTLs displaying trajectories of the mixed type move in an environment where pre-existing channels and therefore deformed collagen matrix exist, leading plausibly to an easier and consequently faster migration even outside the channels, whereas “slow” CTLs move in a local environment where no pre-existing channels exist and average velocity is lower.

The time evolution of the MSD obtained from the model is in a good agreement with the data (exemplary match is shown in Fig. 9). While the model describes the dynamics of T cells very well, the MSD does not contain further distinctive information to differentiate between various migration types of T cells.

**FIG. 9.**
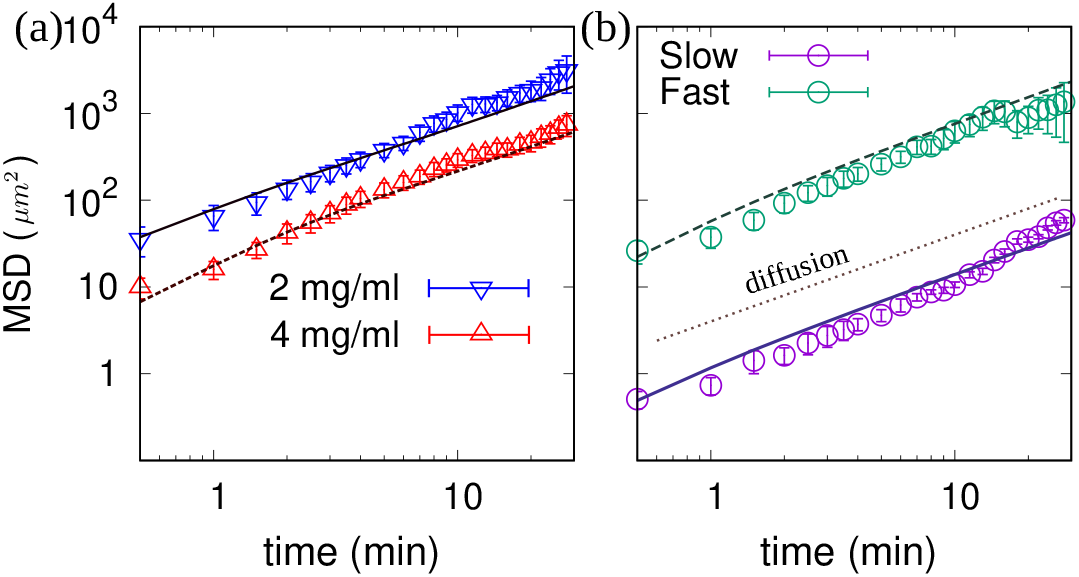
(a) The mean square displacement of mixed motility type in various collagen concentrations obtained from the experiments (symbols) and analytical approach (Eq. 2 with parameters extracted from experimental data summarized in table III) (solid lines) for donor 2.(b) The mean square displacement of slow and fast motility types in collagen concentration 4 mg/ml. The solid lines represent the theoretical estimate of equation 5 withe parameters extracted from the experiments (Table IV).

**TABLE III.** Parameters extracted from the experimental data of mixed type of motility and used in the two-state model.

| density ($\frac{mg}{ml}$) | Donor 1 | | | Donor 2 | | |
| --- | --- | --- | --- | --- | --- | --- |
|  | 2 | 4 | 5 | 2 | 4 | 5 |
| $\langle v \rangle_I (\mu m/s)$ | 0.066 | 0.047 | 0.048 | 0.047 | 0.023 | 0.020 |
| $\langle v \rangle_{II} (\mu m/s)$ | 0.138 | 0.120 | 0.111 | 0.099 | 0.061 | 0.055 |
| $\langle v^2 \rangle_I (\mu m^2/s^2)$ | 0.0189 | 0.0038 | 0.0037 | 0.0149 | 0.0009 | 0.0006 |
| $\langle v^2 \rangle_{II} (\mu m^2/s^2)$ | 0.0510 | 0.0198 | 0.0165 | 0.038 | 0.0055 | 0.0040 |
| $R_I$ | 0.15 | 0.35 | 0.38 | 0.19 | 0.26 | 0.35 |
| $R_{II}$ | 0.62 | 0.59 | 0.59 | 0.33 | 0.56 | 0.69 |
| $\Delta t$ (min) | 0.5 | 0.5 | 0.5 | 0.5 | 0.5 | 0.5 |
| $\kappa_{I \rightarrow II}$ | 0.20 | 0.08 | 0.09 | 0.11 | 0.10 | 0.07 |
| $\kappa_{II \rightarrow I}$ | 0.06 | 0.14 | 0.17 | 0.14 | 0.11 | 0.13 |
| $\langle \tau \rangle_I$ (min) | 2.5 | 6.2 | 5.5 | 4.5 | 5 | 7.1 |
| $\langle \tau \rangle_{II}$ (min) | 8.3 | 3.5 | 2.9 | 3.6 | 4.6 | 3.8 |

**TABLE IV.** Parameters extracted from the experimental data of fast (up) and slow(down) types of motility.

| density ($\frac{\text{mg}}{\text{ml}}$) | Donor 1 | | | Donor 2 | | |
| --- | --- | --- | --- | --- | --- | --- |
|  | 2 | 4 | 5 | 2 | 4 | 5 |
| $\langle v \rangle_F (\mu\text{m/s})$ | 0.17 | 0.12 | 0.09 | 0.11 | 0.07 | 0.07 |
| $\langle v^2 \rangle_F (\mu\text{m}^2/\text{s}^2)$ | 0.033 | 0.019 | 0.013 | 0.018 | 0.008 | 0.006 |
| $R_F$ | 0.54 | 0.41 | 0.45 | 0.53 | 0.40 | 0.56 |
| $\Delta t$ (min) | 0.5 | 0.5 | 0.5 | 0.5 | 0.5 | 0.5 |
| $\langle v \rangle_S (\mu\text{m/s})$ | 0.032 | 0.019 | 0.017 | 0.027 | 0.011 | 0.010 |
| $\langle v^2 \rangle_S (\mu\text{m}^2/\text{s}^2)$ | 0.0014 | 0.0006 | 0.0005 | 0.0010 | 0.0002 | 0.0001 |
| $R_S$ | -0.12 | 0.18 | 0.19 | 0.05 | 0.21 | 0.37 |
| $\Delta t$ (min) | 0.5 | 0.5 | 0.5 | 0.5 | 0.5 | 0.5 |

## V. DISCUSSION

We analyzed the trajectories of CTLs within 3D collagen matrices with different concentrations. We found three motility types in all experiments: slow, fast and mixed. Similar motility types have been reported for natural killer (NK) cells in hydrogel collagen with a concentration of 3 *mg/ml* in the presence of target cells [29]. The similarity of the characteristics of CTLs and NK cell trajectories points towards a common mechanism for migration of both cell types through collagen networks.

A plausible scenario to explain our findings, is that the cells which arrive first in the collagen network perform a persistent random walk unless they move into denser areas of the network, where they become slower, but eventually find a way to move again, which leads to two-state motility. When the cells move through the collagen network, they leave a channel by displacing or stretching collagen fibers. These channels facilitate the movement of other T cells, such that cells entering already existing channels move faster and tend to remain in the existing channel network. They do not switch to slow movement and thus establish the fast type. The slow cells mainly remain in one part of the network and only “wiggle” around and seem to be nearly immobile.

In principle, more and more channels can be built by migrating T cells with time, therefore, leading to an increase in overall migration speed over time. For our experiments, however, the visualization period was about an hour, and we can estimate the amount of new channels that are produced by the T cells: The cell density is 10^7^ cells/ml which gives an average cell-to cell distance of 50 *μ*m. The average velocity of the *slow* cells – which are those that drill *new* channels and are nearly one third of all T cells – is 0.02 *μ*m/sec. If the motion would be totally persistent 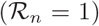 a slow T cell drill within the observation time of 1 hour a 72 *μ*m long new channel, which is not long enough to allow significantly more number of channels to be built. Assuming a cylindrical shape of the channels with a radius of roughly the radius of a T cell, around 5 *μ*m, one can estimate the total volume of new channels created during the observation period in 1 ml to be 0.019 ml, i.e. 2%. of the total volume. This is only a small portion and is the reason why the average velocity does not increase during the observation period. We should also note that before the observation period starts, the T cells were already present for 2 hours in the collagen matrix. This is the reason why right from the beginning of the observation period one observes fast T cells: they move in pre-existing channels (which are rather sparse, though: 11 % vol. We would like to stress that the low channel production rate is the reason why in the theoretical model that we use one can neglect the production of new channels in the observation period: for short observation times the statistics of the cell movement is well reproduced by assuming a static network of channels. Our model includes another simplification: instead of generating and fixing this static network before the observation starts, we assume that it is generated “on the fly” when the cells are in the “fast” mode. This simplification is similar to using the annealed approximation [41, 42] for quenched disorder - which is known to be good for sufficiently high dilution, which is the case for a sparse channel network. Long-term visualizations will be of great help to verify these points. However, as in the current experimental settings, the collagen matrix falls apart in within 8-12 hours depending on the density. Nevertheless, our conclusion is supported by a recent study, which shows that in vivo in salivary gland, T cells follow trajectories established by macrophages, allowing T cells to migrate faster [43].

Compelling evidence shows that when immune cells, including T cells, go through a constricted space, nucleus, as the stiffest organelle in the cells, is the rate-limiting factor [44, 45]. Therefore, when the width of the channels in ECM is smaller than the diameter of nucleus, it would become a speed limiting factor for CTL migration in 3D matrix. If the channels are too narrow for the nucleus to pass, the T cells would appear to halt at the position until the matrix breaks loose or until they manage to squeeze themselves out. According to the experimentally determined pore size distribution in collagen matrices with various concentration [40], one can estimate that the average pore diameter is around 7.5 *μm* for collagen density 2 *mg/ml*. The diameter distribution is broad enoughto have some pores with diameter around 9 *μm*, where T cells with diameter 10 *μm* can squeeze through without deforming the surrounding collagen fibers. On the other hand, in collagen concentrations 4 and 5 *mg/ml*, the average pore diameter is less than 5 *μm* with a very narrow distribution and T cells hardly find pores large enough to squeeze through. This is likely the reason of the different 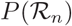 in Fig. 7(a). As a consequence, channel formation of CTLs during the preparation period is even more essential to explain the observed fast migration in denser collagen matrices.

The morphology of the “channel” visible in Fig. 8 is incompatible with a randomly generated filament network (see background). The latter has of course randomly distributed regions with higher and lower filament density but elongated cylindrical tunnels as visible in Fig. 8 with a diameter of approximately equal to the diameter of a T cell and completely void of filaments cannot occur by chance with significant probability. The cylindrical tunnels could be produced by T cells, as observed in Suppl. Movie 4, as they pass through, and if some collagen fibers do not completely spring back to their original position, leaving a broadened channel behind. Another possibility is that these channels have been produced by either T cells degrading the local matrix by secretion of matrix metalloproteases (MMP) or by T cells tearing matrix apart by exertion of mechanical forces during the 2 hours before the observation and tracking was started - and leaving behind elongated, cylindrical tunnels of approximately the same diameter as T cells. Concerning the former option, it is reported that treatment of MMP inhibitors in human CD4 T cell blasts does not affect T cell speed [46]. In CD4^+^ T cells, MMP2 and MMP9 are expressed [47], which do not degrade collagen type I [48], which was used in our experiments. The collagen type I-degrading MMPs (MMP1, MMP8, MMP12 and MMP14) are not expressed in bead-stimulated primary human CD8^+^ T cells [49]. Due to lack of MMPs, channel formation most probably proceeds via collagen filament deformation or destruction rather than degradation.

In summary, the aim of the present study was to analyze the migration dynamics of CTLs in collagen matrices with different densities to understand potential differences in T cell migration patterns and to elucidate the role of collagen density. The investigation of the effects of the migration patterns that we reveal for the search efficiency of CTLs remains for future studies. Based on our observations, we expect that the search efficiency of T cells decreases in dense collagen matrices because even though the migration pattern per se does not differ, furthermore the velocity of CTLs decreases in dense ECM. Interestingly, T cells “dig” into the collagen network and create channels that are later used by other T cells. This is beneficial for search optimization as T cells can move faster in the channels and fast migration results in reduced search times when searching for static or dynamic obstacles.

## VI. APPENDIX

We confine ourselves to a two-dimensional model to derive an analytical formula for the mean square displacement of one Cartesian coordinate 〈*x*^2^(*t*)〉, which is then multiplied by 3 to give the prediction for the MSD in three dimensions 〈*r*^2^(*t*)〉 = 3〈*x*^2^(*t*)〉[50]. A Fourier-*z*-transform technique [51] was employed to solve the master equations 1. The *z*-transform *A*(*z*) of an arbitrary function *A_n_* of a discrete variable *n*=0, 1, 2, … is defined as 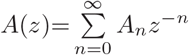, which is equivalent to Laplace transform in a continuous-time description. The exact result for the mean square displacement is obtained via the inverse *z*-transform of the following equation [31]:

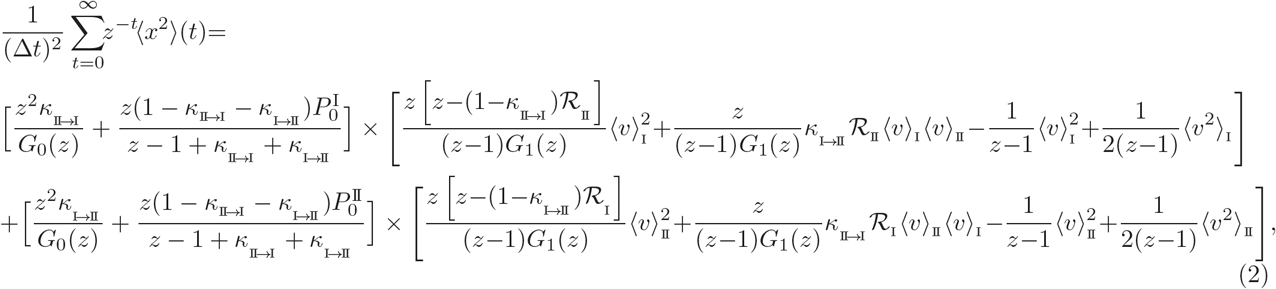

with

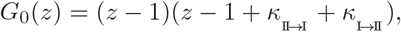

and

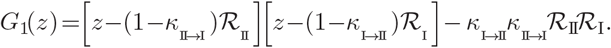

In equation 2, 〈*v*〉_I_, 〈*v*〉_II_, 〈*v*^2^〉_I_ and 〈*v*^2^〉_II_ are first and second moments of velocity of T cells in states I and II. 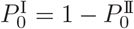 is the initial condition and shows the probability of starting the motion in state I or equivalently the percentage of all T cells in state I at the beginning of tracking. The initial condition 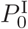 only affects the short-time behavior of motion, we set this parameter to 0 for all cases. This means that we assume all T cells start their motion in the faster mode. 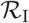 and 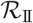 are the Fourier transform of distributions of turning angle *R_I_* (*ϕ*) and *R_II_* (*ϕ*) in Eqs. 1:

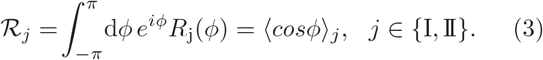

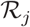 varies from −1 for reversing the direction (i.e. *ϕ*=±*π*) to 0 for a uniformly random turning (*ϕ*∈[−*π, π*]) and to 1 for continuing along the previous direction of motion (i.e. *ϕ*=0).

The master equation for the probability distribution 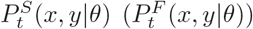 describing the slow (fast) motility type is identical to the master equation for 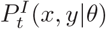 with the switching rate *κ*_*I*→*II*_ (*κ*_*II*→*I*_) set to zero (see Eq.1):

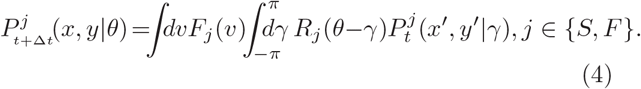

where 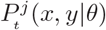 is the probability density of a T cell to arrive at position (*x, y*) with direction *θ* at time *t* and *F_j_*(*v*) and *R_j_*(*θ*) are the distribution functions of velocity and turning angle, respectively. The resulting MSD in this case will be:

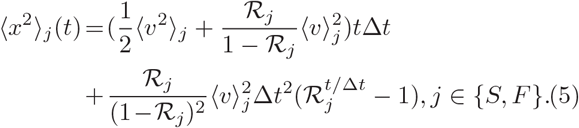

For large times the term proportional to *t* dominates the r.h.s. (since 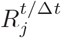 vanishes for *t* → ∞, which implies conventional diffusive behavior. For short times *t*−*t*_0_ → 0 the MSD depends algebraically on *t*, 〈*x*^2^〉 ~ *t*^*α*^ with an exponent given by [Shaebani14]:

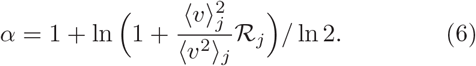

For ballistic motion, i.e. 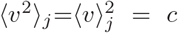, 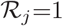, one obtains *α* = 2, and for conventional diffusion, 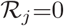, one obtains *α* = 1.

## SUPPLEMENTARY INFORMATION

Supplementary information accompanies this paper, including four movies.

## AUTHOR CONTRIBUTIONS

Z.S., B.Q., M.H. and H.R. designed the research. R.Z. and B.Q. performed the experiments. Z.S. analyzed the experimental data and employed the analytical model. All authors contributed to the interpretation of the results. Z.S., B.Q. and H.R. wrote the manuscript which was edited by all authors. Correspondence should be addressed to Z.S.

## ACKNOWLEDGEMENTS

We acknowledge financial support from Collaborative Research Center SFB 1027, M.H. from BMBF grant 031L0133 and R.Z. from HOMFOR2018 grant.

